# A peptidergic amygdala microcircuit modulates sexually dimorphic contextual fear

**DOI:** 10.1101/2020.01.28.923482

**Authors:** AK Rajbhandari, JC Octeau, S Gonzalez, ZT Pennington, J Trott, J Chavez, E Ngyuen, N Keces, WZ Hong, RL Neve, J Waschek, BS Khakh, MS Fanselow

## Abstract

Trauma can cause dysfunctional fear regulation leading some to develop disorders like post-traumatic stress disorder (PTSD). The amygdala regulates fear, and, PACAP and PAC1 receptors are linked to PTSD symptom severity at genetic/epigenetic levels, with a strong link in females with PTSD. We discovered a PACAPergic projection from the basomedial amygdala (BMA) to the medial intercalated cells (mICCs). In vivo optogenetic stimulation of this pathway increased cfos expression in mICCs, decreased fear retention and increased fear extinction. Selective deletion of PAC1 receptors from the mICCs in females reduced fear acquisition, but enhanced fear generalization and reduced fear extinction in males. Optogenetic stimulation of the BMA-mICCs PACAPergic pathway produced excitatory postsynaptic currents (EPSCs) in mICCs neurons, which was enhanced by PAC1 receptor antagonist, PACAP 6-38. Our findings show that mICCs modulate contextual fear in a dynamic and sex-dependent manner via the microcircuit containing the BMA and mICCs, dependent on behavioral state.

## Introduction

Given the very high prevalence of anxiety-related pathology in society, it is essential that we understand the cellular and circuit mechanisms underlying emotion dysregulation [1]. The amygdala and its associated structures play a key role in processing and reacting to emotional stimuli and it is known to be involved in anxiety disorders such as Post Traumatic Stress Disorder (PTSD) [2]. The cortex-like regions of the amygdala proper (lateral and basal nuclei of the basolateral amygdala complex-BLA) receive sensory information from neocortex and thalamus [3]. Plasticity within these nuclei supports associative learning of sensory information pertinent to affect and supports processes such as Pavlovian fear conditioning [4-6]. On the other hand, generation of most fear-related behaviors is initiated by the nearby medial portion of the striatal-like Central Nucleus (CN, Swanson & Petrovich, 1998) [3]. There are several routes of communication between the amygdala and CN, only some of which are direct [7]. Indirect microcircuits include relays in the lateral portion of the CN [8, 9] as well as clusters of GABAergic cells that lie in the capsule separating amygdala and CN [7]. There is currently only limited information about which of the specific aspects of fear learning are selectively served by these separable microcircuits. The capsular, or medial intercalated cell clusters (mICCs), appear to play a role in fear extinction. A majority of the mICCs express mu-opioid receptors on their cell bodies and selective ablation of these neurons partially eliminates recall of fear extinction, the loss of fear responses to a stimulus that previously triggered fear due to repeated exposure without any aversive consequences [10]. Whether or not the mICCs are involved in other aspects of fear, such as acquisition, recall, and generalization is unknown. Therefore, the present experiments combined a contextual fear conditioning task that allowed us to interrogate each of these behaviors with a novel approach to target the medial ICCs and to dissect specific neuronal pathways expressed in this circuitry.

The BLA complex contains both excitatory and inhibitory neurons and the neurotransmitters and neuromodulators it utilizes to communicate with the mICCs are largely unknown. Pituitary adenylyl cyclase-activating peptide (PACAP) and its G-protein coupled receptor, PAC1, are expressed in several brain areas involved in emotion and arousal including the amygdala and mICCs. PACAP/PAC1 have been linked to PTSD symptom severity at both genetic and epigenetic levels, and the genetic link is especially strong in females with PTSD [11]. Using several lines of mice that genetically targeted PACAP or PAC1 expressing cells, we identified a microcircuit consisting of PACAP expressing neurons in the basal medial nucleus of the amygdala (BMA) that project to PACAP/PAC1 expressing mICCs and discovered that this microcircuit regulates fear acquisition, generalization, recall and extinction in a sex-dependent manner.

## Results

### 1. PACAP-expressing neurons in the BMA innervate the medial ICCs

We examined EGFP expression in the Adcyap1-EGFP mice, which restricts EGFP expression to PACAP expressing neurons [12]. Although distributed broadly in the amygdala, EGFP^+^ cells were enriched in the lateral and basomedial nucleus (BMA) subregion with fibers in the mICCs (Fig. 1A and Fig. S1A, B). We evaluated the local afferents of BMA and found a majority of PACAPergic neurons in the BMA innervate the dorsal and ventral mICCs (Fig.1A). By comparison, we found little to no innervation of the CN by EGFP^+^ neurons (Fig 1A. and Fig. S1B). The pattern of expression corresponds with PACAP mRNA expression, which is high in the BMA and with some expression in the mICCS (Allen Brain Atlas Mouse Brain *In Situ* hybridization) (Fig. S1C). To further confirm the existence of monosynaptic PACAPergic projections from BMA to mICC we used an intersectional approach as described in Fenno et. al. 2014 [13]. For this, we injected a retrogradely trafficked Cre-dependent HSV virus (hEF1α-LS1L-mCherry-IRES-flpo) into the mICCs (dorsal portion) and a Flp-dependent AAV5-EF1a-fDIO-ChR2-eYFP-WPRE into the BMA of Adcyap1-2A-Cre. Adcyap1-2A-Cre mice express Cre specifically in PACAP-containing neurons. With the intersectional approach, the HSV virus, which is Cre-dependent and expresses FLP, retrogradely transports allowing expression specifically in PACAPergic neurons in the BMA. The AAV viruses in turn are FLP-dependent, so therefore expresses only in neurons that have FLP. This allows labeling specific projections from the BMA to mICCs. Thus, using two different approaches, we were able to confirm that PACAPergic (Adcyap1-EGFP) neurons in the BMA innervate mICCs (Fig. 1B and Fic. 1C).

**Fig.1.**
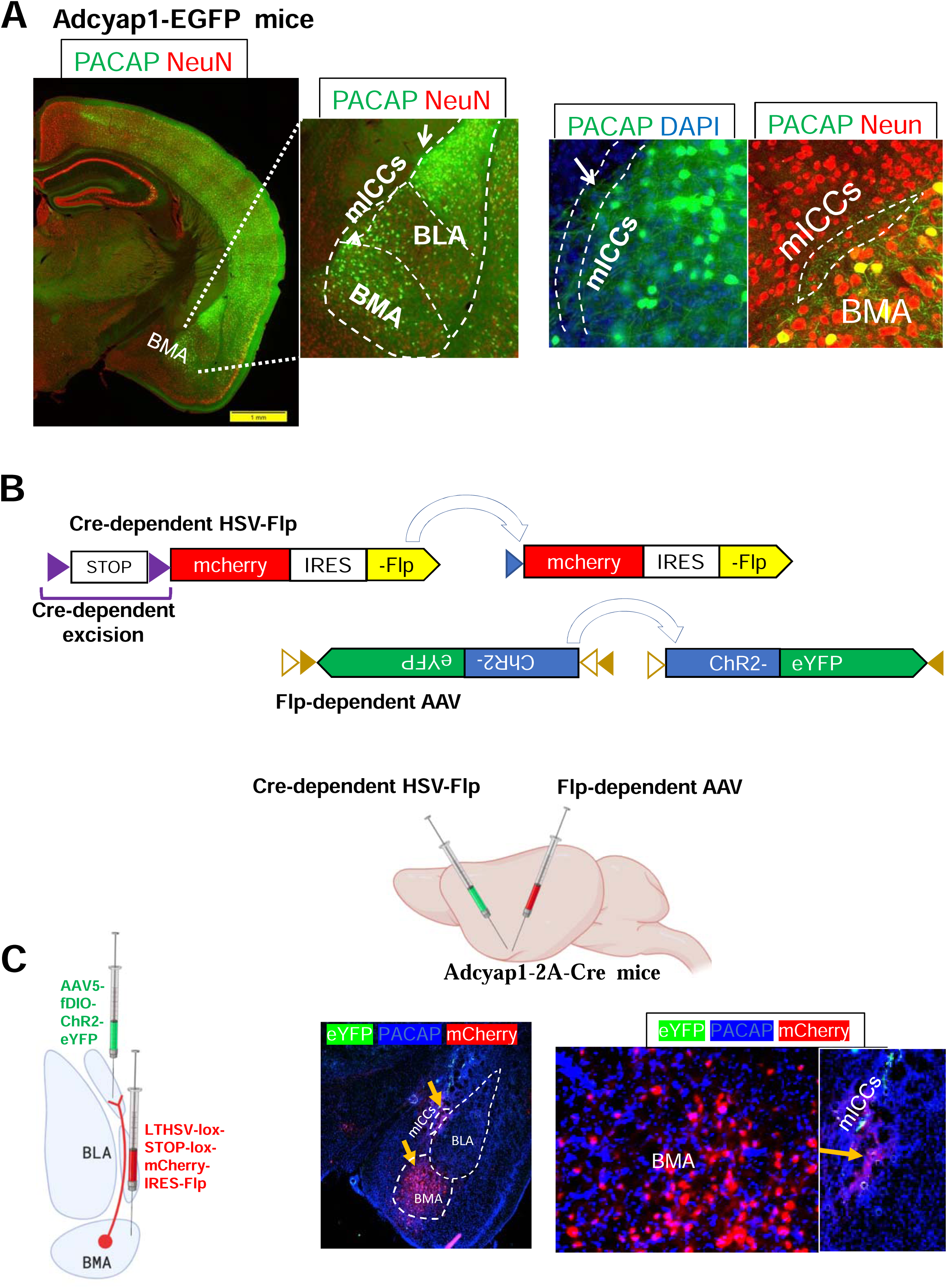
PACAP-expressing neurons in the BMA innervate the medial ICCs. **A.** Representative images showing immunohistochemistry for green fluorescent protein (GFP) in Adcyap1-EGFP mice. Left: PACAPergic neurons are highly expressed in the BMA and terminals of those neurons are innervate the mICCs and CeA. Right: Panels show areas at 40× with PACAPergic innervation in the mICCs. **B.** Images depicting the viral constructs (top) for the intersectional viral injections in Adcyap1-2A-Cre mice (bottom). The Cre-dependent LTHSV-lox-STOP-lox-mCherry-IRES-Flp virus that was injected in the mICCs is shown on the top and the AAV5-fDIO-ChR2-eYFP that was injected into the BMA is shown in the bottom. **C.** Left: Cartoon depicting the injection strategy for the intersectional approach in Adcyap1-2A-Cre mice. LTHSV-lox-STOP-lox-mCherry-IRES-Flp virus was injected into the dorsal mICCs and AAV5-fDIO-ChR2-eYFP was injected into the BMA. Right: Representative image panels showing expression of ChR2 in PACAPergic neurons in the BMA innervating the mICCs.

### 2. *In vivo* optogenetic stimulation of BMA PACAPergic input to the mICCs decreases fear retention and increases fear extinction

Next we wished to determine if, and how, this BMA (PACAP) to mICCs (PAC1) pathway contributes to the learning and expression of fear. Using the same intersectional approach in the Adcyap1-2A-Cre mice, we conducted an optogenetic gain of function experiment. We carried out bilateral *in vivo* optogenetic stimulation of PACAPergic fibers in the mICCs that emanate from the BMA and tested different aspects of contextual fear behavior including acquisition, generalization, retention and extinction. The behavioral procedure is shown in Fig. 2A. These experiments were carried out in both males (N=8 (ChR2); N=9 (EYFP)) and females (N=9 (ChR2); N=7 (EYFP)). Acquisition measures the ability of animals to learn the association between the context and shock and an asymptotic level of conditional fear is graded in a manner that is proportional to shock intensity. ANOVA revealed that there was no Sex X Group interaction in fear acquisition indicating that the animals acquired/learned fear in a similar manner (F=0.833; p>0.05) (Fig. 2C). Learned responses generalize to other contexts and overgeneralization occurs in anxiety disorders reflecting fear in an inappropriate context. Moderate levels of freezing in this alternate context indicate that the mice exhibit over-generalized fear. However, we found that the Sex X Group interaction in the generalization test was not significant, indicating that the animals did not differentially generalize fear to a novel context (Fig 2D) (F=0.594; p>0.05).

**Fig. 2.**
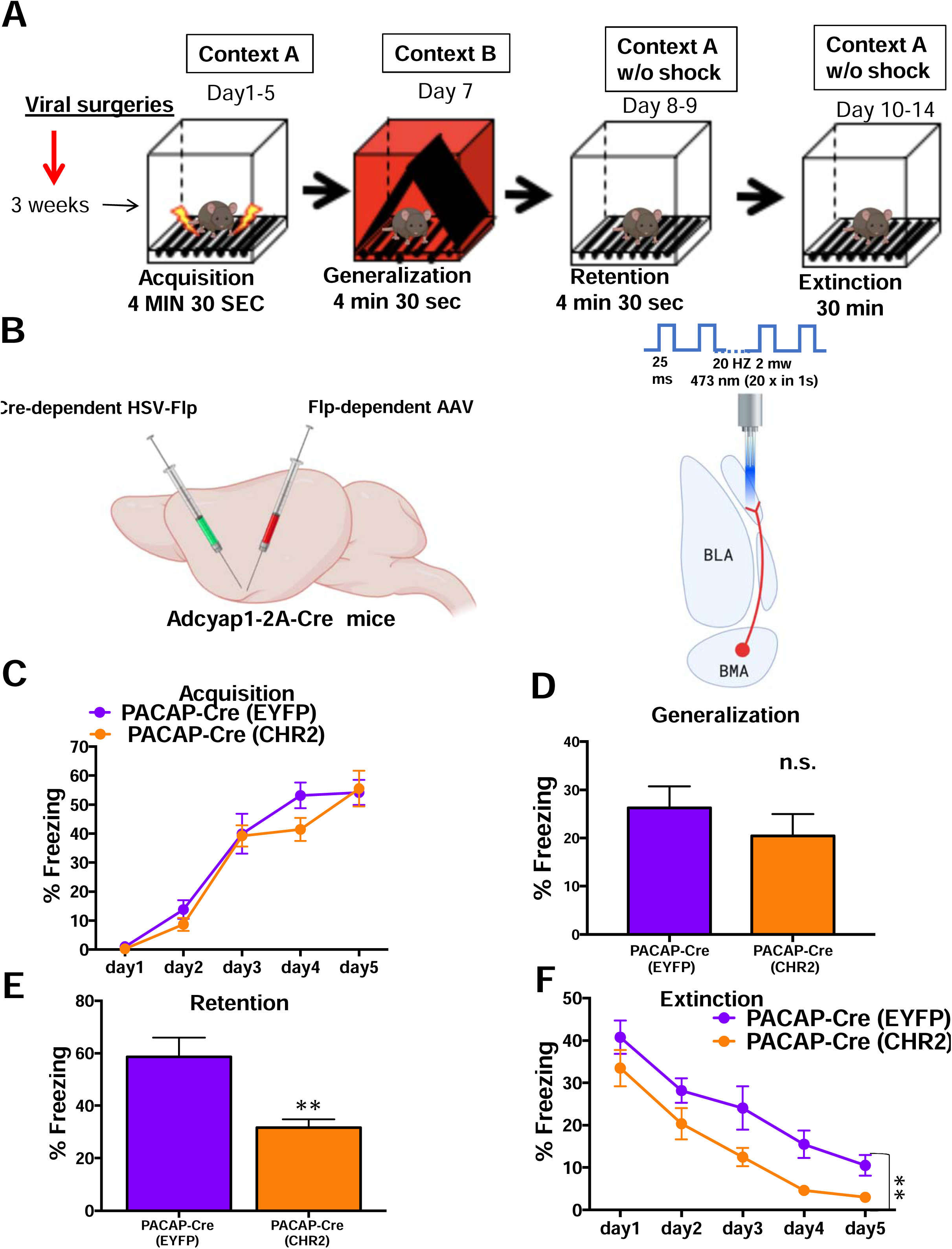
*In vivo* optogenetic stimulation of PACAPergic inputs to the mICCs from BMA decreases fear retention and increases fear extinction. **A.** Cartoon diagram depicting viral injection surgeries timeline and behavior protocol. Mice went through fear acquisition in context A, generalization test in context B, retention test in context A and extinction in context A. **B.** Left: Cartoon depicting the intersectional viral injection strategy in the BMA and mICCs. Right: Cartoon depicting the optogenetic stimulation strategy and parameters for stimulation. Males (N=8 (ChR2); N=9 (EYFP)) and females (N=9 (ChR2); N=7 (EYFP)) were used in these experiments. **C.** Graph showing freezing during acquisition. There was no significant Sex X Group interaction in fear acquisition (F=0.833; p>0.05). However, we found that the. **D.** Graph showing freezing during fear generalization test. The Sex X Group interaction in the generalization test was not significant, indicating that the animals did not differentially generalize fear to a novel context (F=0.594; p>0.05). **E.** Graph showing freezing during fear retention test. There was a significant main effect of Group (F=13.08; p<0.05). Post-hoc comparison indicated that the group that received optogenetic stimulation of the PACAPergic neurons from BMA to mICCs showed a significantly reduced level of freezing compared to the controls. However, there was no significant Group X Sex interaction (F=0.008; p>0.05). **F.** Graph showing freezing during extinction. There was a main effect of Group during the extinction test (F=20.128; p<0.05), but no Group X Sex interaction (F=0.1; p>0.05).

The retention test was designed to measure the ability of animals to maintain a long-term fear memory of the context in which they acquired fear. We found a significant main effect of Group (F=13.08; p<0.05) in the retention test, such that the group that received optogenetic stimulation of the PACAPergic neurons from BMA to mICCs showed a significantly reduced level of freezing compared to the EGFP controls (Fig. 2E). However, there was no significant Group X Sex interaction (Fig 2E) (F=0.008; p>0.05).

Extinction is the loss of expression of learned behavior with repeated exposure to the conditional stimulus (e.g. context) without the unconditional stimulus (e.g. shock). We found a main effect of Group during the extinction test (F=20.128; p<0.05), but no Group X Sex interaction (Fig 2F) (F=0.1; p>0.05). These results show that stimulation of PACAPergic pathway from BMA to mICCs alters specific aspects of fear behaviors. Specifically, our results indicate the activation of PACAPergic pathway from BMA to mICCs decreases retention of fear indicating that these animals had a weaker memory of the context where they acquired fear behaviors. These animals also showed enhanced extinction of fear, indicating that activation of PACAPergic pathway from BMA to mICCs reduces fear of the context in which the context and shock associations were learned.

### 3. Deletion of PAC1 receptors from the mICCs enhances fear generalization and decreases fear extinction in males but not in females

Previous studies have shown that forebrain-specific deletion of PAC1 receptors leads to an impairment of contextual fear conditioning, but in these studies other aspects of contextual fear learning such as generalization and extinction were not examined [14]. *In situ* hybridization shows that PAC1 receptor mRNA expression is high in the mICCs compared to BLA/BMA or CN (Allen brain Atlas and Fig. S2B). We conducted a loss of function experiment to determine how a corresponding loss of mICCs PAC1 receptor gene expression affects the same set of fear behaviors as we tested with the optogenetic gain of function experiments. While mICCs are difficult to target because of their small size, we were able to localize our viral injections to these cells (Fig. S2C and Fig. S2D). Only the animals that had bilateral and specific viral expression localized in the mICCs were used in the behavioral analysis. We carried out testing in both males (N=14 (Cre); N=16(GFP) and females (N=15 (Cre); N=18 (GFP). ANOVA revealed a main effect of Day (F=201.186, P<0.05), Sex (F=10.586, p<0.05) and a Sex X Group interaction (F=4.757, p<0.05) in acquisition (Fig. 3B and 3E). There was a main effect of Group (F=3.858, P=0.05) and a Sex x Group interaction (F=9.352, P<0.05) on generalization test (Fig. 3C and 3F). Post-hoc analysis showed that males with PAC1 deletion were significantly different from control (P<0.05). There was no effect of Sex or Group or Group x Sex interaction on the retention test (Fig. S3D) (F=2.463; p>0.05). There was a main effect of Day (F=49.711, P<0.05), Sex (F=12.513, p<0.05) and Day x Sex x Group interaction (F=2.545, p<0.05) in extinction. Post-hoc analysis revealed that extinction rate in males with PAC1 deletion was significantly higher than controls on days 2, 3 and 4 of extinction (P<0.05). Females showed decreased fear acquisition, but males showed enhanced fear generalization and decreased fear extinction (Fig. 3C-Fig.3H). We verified that viral expression was localized in the mICCs with post-mortem analysis using the RNAscope technique; PAC1 mRNA in the mICCs was significantly reduced after injection of AAV-Cre compared to AAV-GFP control (P<0.05; Fig. 3H).

**Fig. 3.**
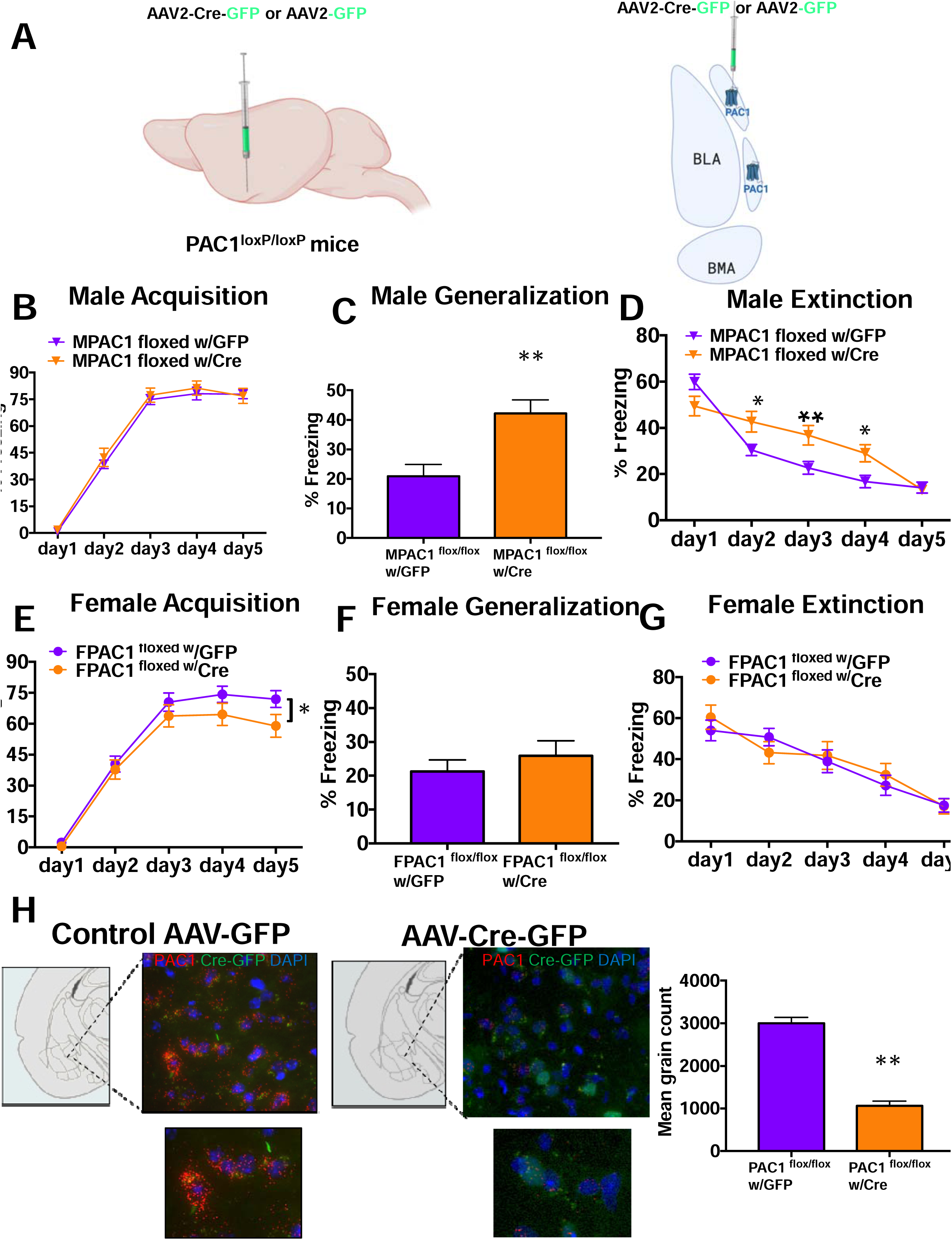
Deletion of PAC1 receptors from the mICCs of PAC1 ^loxp/loxp^ mice enhances fear generalization and decreases fear extinction in males but not in female. **A.** Images depicting the virus injection strategy on the left and right in the mICCs. Males (N=14 (Cre); N=16(GFP) and females (N=15 (Cre); N=18 (GFP) were used in these experiments. **B.** and **E**. Graphs showing freezing during acquisition in males (top panel) and females (bottom panel). ANOVA revealed a main effect of Day (F=201.186, P<0.05), Sex (F=10.586, p<0.05) and a Sex X Group interaction (F=4.757, p<0.05) in acquisition. **C.** and **F**. Graphs showing freezing during fear generalization test. There was a main effect of Group (F=3.858, P=0.05) and a Sex x Group interaction (F=9.352, P<0.05) on generalization test. Post-hoc analysis showed that males with PAC1 deletion were significantly different from control (P<0.05). **D.** and **G**. Graph showing freezing during extinction. There was a main effect of Day (F=49.711, P<0.05), Sex (F=12.513, p<0.05) and Day x Sex x Group interaction (F=2.545, p<0.05) in extinction. Post-hoc analysis revealed that extinction rate in males with PAC1 deletion was significantly higher than controls on days 2, 3 and 4 of extinction (P<0.05). **H.** Example panels showing PAC1 expression levels in the mICCs from mice with AAV-GFP and AAV-Cre-GFP using the RNAscope in situ hybridization technique. Right panel: Graph showing mean grain count of PAC1 mRNA PAC1 mRNA in the mICCs was significantly reduced after injection of AAV-Cre compared to AAV-GFP control (P<0.05).

### 4. *In vivo* optogenetic activation of PACAP-expressing neurons in the BMA enhances expression of cfos in the mICCs

Next we wanted to determine if activation of the BMA PACAPergic neurons would alter cellular activity of mICCs neurons. Using the same intersectional approach in the Adcyap1-2A-Cre mice, as described previously, we performed *in vivo* optogenetic stimulation of the BMA in anesthetized mice and measured changes in expression of cfos in the mICCs. The intersectional Cre-dependent HSV-Flp and Flp-dependent AAV viruses were injected into the mICCs and BMA respectively in both hemispheres. The optic fiber was also implanted in both hemispheres targeting the BMA (Fig. 4A, B and C). We performed optogenetic stimulation of BMA PACAPergic neurons in only one hemisphere (Fig. 4B). This analysis revealed that the number of cfos+ cells was significantly enhanced in the mICCs in the hemisphere with optogenetic stimulation of chR2-contaning fibers of the BMA. Paired T-Test showed that optogenetic stimulation of BMA significantly enhanced expression of cfos in the mICCs in the hemisphere with optogenetic stimulation compared to the mICCs of the hemisphere without optogenetic stimulation (N=4 (2M;2F), df=3, p<0.001) (Fig. 4D and Fig. 4E). This result shows that altering activity of PACAPergic neurons in the BMA modulates activity of neurons in the mICCs.

**Fig. 4:**
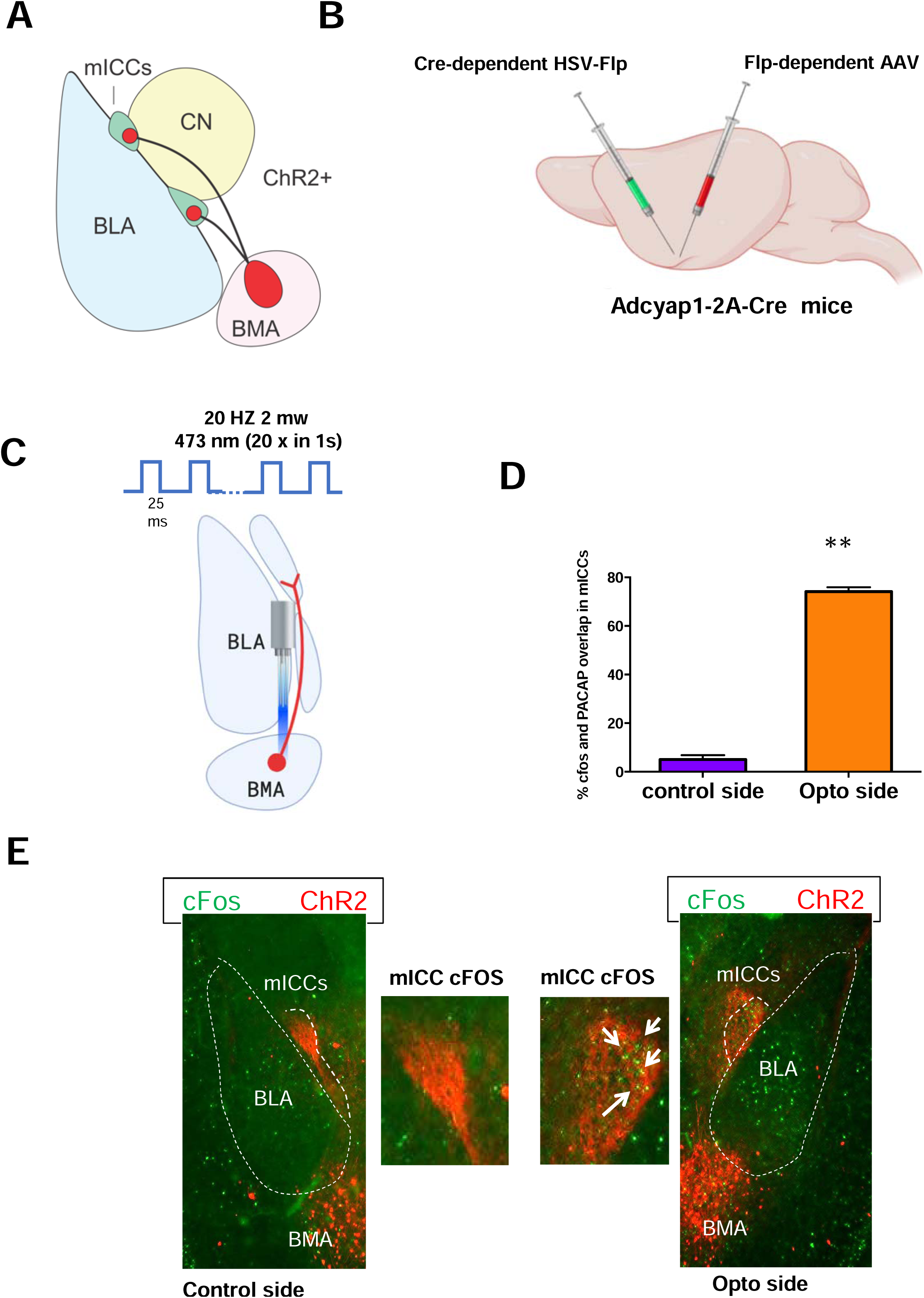
*In vivo* optogenetic activation of PACAP-expressing neurons in the BMA enhances expression of cfos in the mICCs. **A.** Cartoon depicting the location of the mICCs and BMA relative to central (CeA) and basolateral (BLA) nuclei of the Amygdala. ChR2-expressing cells within the BMA are depicted in red. **B.** Cartoon depicting optogenetic stimulation strategy for stimulating the BMA in anesthetized Adcyap1-2A-Cre mice. The Cre-dependent LTHSV-lox-STOP-lox-mCherry-IRES-Flp virus that was injected in the mICCs and the AAV5-fDIO-ChR2-eYFP that was injected into the BMA. **C.** Optogenetic stimulation strategy for stimulating the BMA neurons that innervate mICCs. **D.** Graph showing expression level of cfos that was co-localized with mcherry in dorsal the mICCs was significantly enhanced in the hemisphere where optogenetic stimulation was delivered. **E.** Representative image panels with immunohistochemistry for cfos (green) and mcherry (red) expression showing enhanced cfos expression in the mICCs hemisphere where optogenetic stimulation was carried out in the BMA.

### 5. *Ex vivo* optogenetic stimulation of the BMA-ICC PACAPergic pathway enhances EPSCs in the mICCs that is further enhanced by application of a PAC1 receptor antagonist

Next, we wanted to determine if stimulation of BMA PACAPergic terminals changes synaptic activity of mICC neurons. For this, we performed *ex vivo* electrophysiological recordings in combination with optogenetic stimulation of BMA PACAPergic fibers expressing ChR2. We injected AAV5-EF1a-DIO-hChR2(H134R)-mCherry virus into the BMA of Adcyap1-2A-Cre mice, which expresses Cre in PACAP-containing neurons (both males and females). This allows expression of ChR2 in the efferent pathways of the BMA containing PACAP. We then performed whole-cell patch clamp recordings from the dorsal mICC region (Fig. 5B). The mICC neurons showed a sagging current upon hyperpolarization and action potentials in response to stepwise changes in current (Fig. 5B). For all electrophysiological experiments, we confirmed that the recording sites were within the mICCs by filling the recorded neurons with biocytin and confirming the appropriate location of the neurons post-hoc (Fig. 5C and Fig.S5). The electrophysiological and morphological properties of mICC neurons in our studies matched previously described properties of mICC neurons[15-17].

**Fig. 5:**
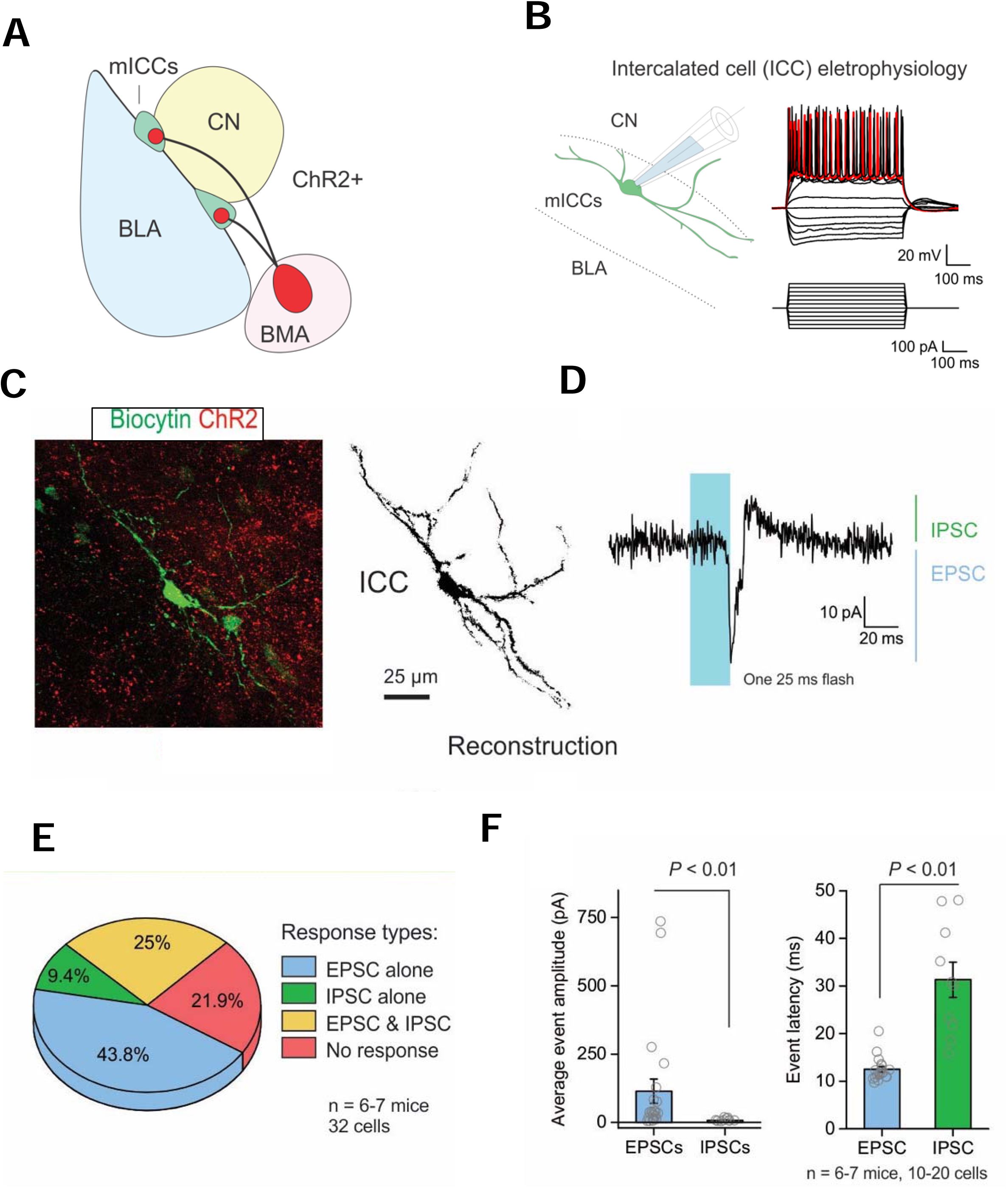
*Ex vivo* optogenetic stimulation of BMA-ICCs PACAPergic pathway enhances EPSCs in the mICCs. **A.** Cartoon depicting the location of the ChR2-expressing cells within the BMA in red and mICCs in green. **B.** *Left:* Cartoon depicting the whole-cell patch clamp of a neuron in the mICC nucleus. *Right:* Whole-cell electrophysiological properties of ICCs with the current injection parameters depicted below. **C.** Image of a biocytin filled mICC neuron (left) and reconstruction (right). **D.** Example trace depicting the electrophysiological responses of neurons from the mICCs relative to the LED flash. In this case, the neuronal response comprises of a EPSC followed by an IPSC response. **E.** Pie chart depicting the classification of response types from ICCs. Majority of mICCs neurons showed EPSCs. **F.** Left: Graphs depicting the amplitudes of EPSC (*left*) or IPSC (*right*) responses of neurons from the mICCs. Right panel: Graph depicting the response latency of either EPSC or IPSC events in mICCs in response to ChR2 activation of PACAP projections.

Optogenetic stimulation of PACAPergic neurons from BMA was carried out with a 473nm light pulsed with a 20 Hz train of 25 ms single pulse or multiple pulses using an LED. A single flash produced excitatory postsynaptic currents (EPSCs) in a majority of mICC neurons (Fig. 5D and 5E). In some cases, EPSCs were followed by inhibitory post-synaptic potentials (IPSCs) (Fig. 5D and 5E). Out of 32 recorded neurons, 43.8% showed EPSCs, 9.4% showed IPSCs, 25% showed EPSCs followed by IPSCs and 21.9% of cells showed no response (N=8). Mann-Whitney U test showed that the average amplitude of EPSCs was enhanced compared to IPSCs (P<0.01; Fig. 5F). For the neurons that showed EPSCs followed by IPSCs, Mann-Whitney U test showed that the IPSC event latencies were much longer compared to the EPSCs (P<0.01; Fig. 5F). Paired sample signed test showed that optogenetic light evoked EPSCs and IPSCs were blocked by the application of CNQX in both cases indicating that the BMA neurons release glutamate (N=6; 11 cells (EPSCs); 6 cells (IPSCs); P<0.01 both; Fig. 4A). Application of the PAC1 receptor antagonist peptide, PACAP 6-38 in the bath significantly enhanced EPSCs (N=6; 14 cells; P<0.05; Fig. 6B). However, application of PACAP 6-38 did not alter paired-pulse ratio (N=6; 14 cells; Fig. 6B).

**Fig 6:**
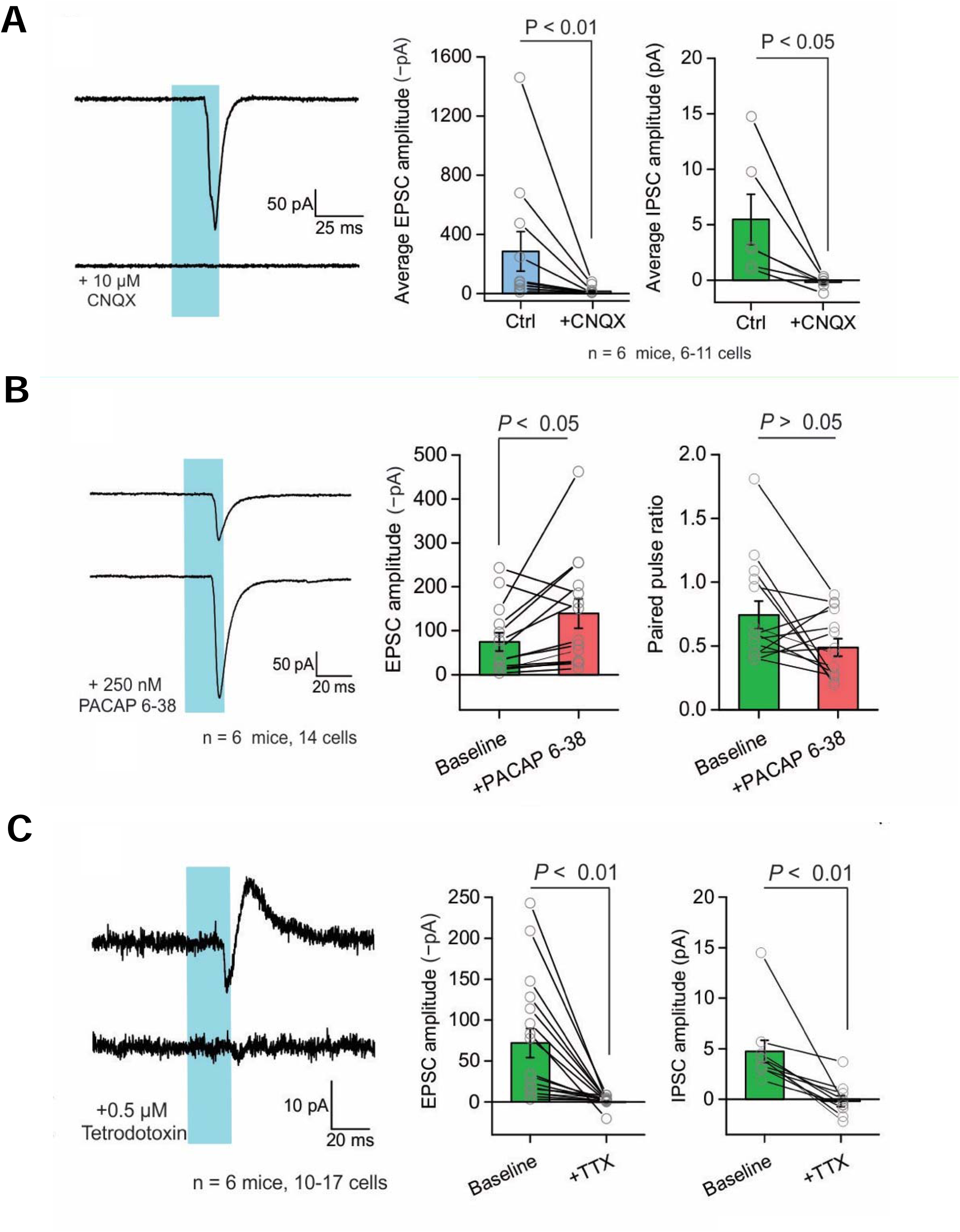
Application of PAC1 antagonist PACAP 6-38 enhanced EPSCs in the mICCs after stimulation of BMA-ICCs PACAPergic pathway. **A.** *Left panel:* Representative current trace from mICC neuron in response to single flashes with or without CNQX applied. Average EPSC (*left*) or IPSC (*right*) amplitude in response to single flashes with or without CNQX. **B.** *Left panel:* Representative current trace from mICC neuron in response to single flashes with or without PACAP 6-38 applied. Average EPSC amplitude (*middle*) or paired-pulse ratio (*right*) in response to single flashes with or without PACAP 6-38. **C.** Representative current trace (*left*) from mICC neuron in response to single flashes with or without TTX applied. Average EPSC (*middle*) or IPSC (*right*) amplitude in response to single flashes with or without TTX.

**Fig. 7.**
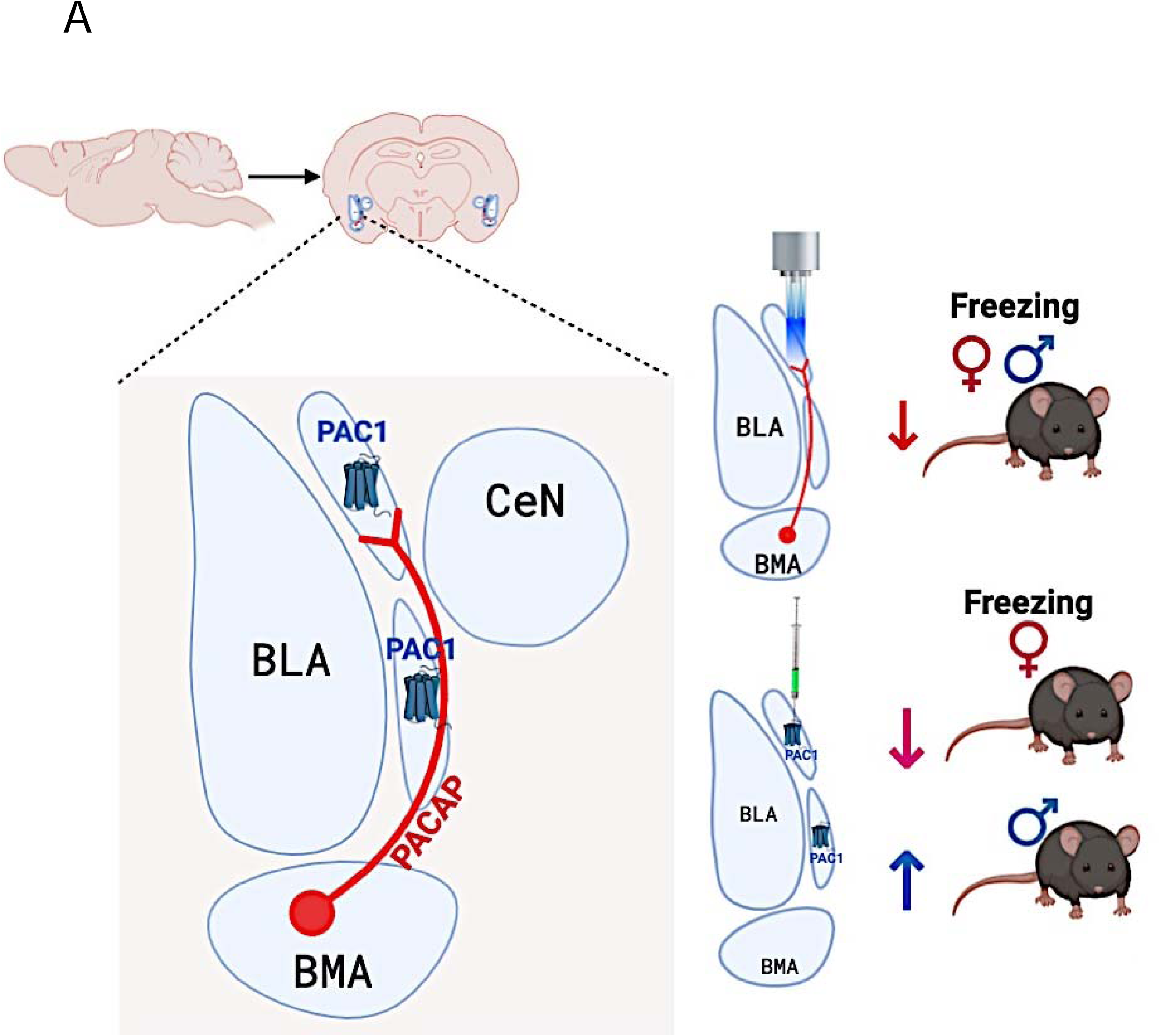
Working model of PACAP in the BMA and PAC1 in the mICCs in fear regulation through amygdala microcircuitry. **A.** A representative model of the amygdala microcircuitry containing PACAP neurons and PAC1 receptors in the BMA and mICCs. i) Stimulating PACAPergic terminals in the mICCs decreased freezing, specifically in the generalization test and extinction. ii) Deletion of PAC1 receptors from mICCs in the females decreased fear acquisition, but deletion of these receptors from mICCs of males enhanced fear generalization and decreased fear extinction.

Application of TTX completely abolished EPSCs and IPSCs (P<0.01; Fig 6C). With bursts of optogenetic stimulation, EPSCs amplitude decreased with an increase in stimulus number (Fig. S4A; N=6; 17 cells (EPSCs); 10 cells (IPSCs)). The IPSCs that were observed in some neurons after EPSCs, were blocked by application of bicuculline (N=6; 11 cells; P<0.05; Fig. S4B). In our studies, we did not find any significant differences in the electrophysiological properties between males and females and therefore the data for males and females was combined. These results indicate that PACAP neurons produce a predominantly excitatory glutamatergic influence on mICCs, which in turn triggers local GABAergic inhibition between mICCs. Overall, these results show that the PACAPergic neurons from BMA to mICCs are synaptically coupled. These results also suggest that PACAP could be co-released with glutamate from BMA neurons mitigates the post-synaptic influence of glutamate.

## Discussion

We describe a previously uncharacterized amygdala microcircuit containing the neuropeptide PACAP and PAC1 receptors in contextual fear regulation, whereby BMA PACAPergic neurons modulate diverse aspects of fear behavior potentially through PAC1 receptors in the mICCs. PAC1 receptor containing neurons in the mICCs, in turn, regulate fear in a sexually dimorphic manner, potentially by differentially recruiting downstream projections to CN or through topographically distinct microcircuit in males and females. We showed our results through gain of function optogenetics, loss of function experiments using genetically modified mice, and electrophysiology. Thus, BMA to mICCs projecting PACAPergic neurons and PAC1 receptors in the mICCs are important nodes in the fear circuitry that control heterogeneous aspects of contextual fear. While mICCs have previously been shown to regulate fear extinction, our studies for the first time demonstrate a new set of roles for the mICCs. Besides fear extinction, the mICCs dynamically regulate several components of fear memory including acquisition, generalization and recall.

Optogenetic stimulation of the PACAP BMA-mICCs pathway decreased contextual fear retention and reduced the expression of fear during extinction. We did not find differences in fear acquisition or generalization. These results suggest that activation of this circuit diminishes expression of acquired fear. Activation of this specific neural circuitry may promote decreased fear by gating the ability for basolateral amygdala complex neurons to promote behavior driving output of the CN or by directly influencing activity of CN output neurons. Studies do show that long term depression occurs in dorsal ICCs after a mild fear stimulus [18]. In our studies, the viral placements were mostly in the dorsal portion of the mICCs. Studies have shown differential expression of Zif, an immediate early gene is different in dorsal and ventral mICCs after fear expression versus fear extinction [19]. It is possible that different sub-divisions of the mICCs modulate different aspects of fear via specific neuropeptide/neurotransmitter and also related to the sex-differences observed in our studies [19, 20]. Interestingly though, morphological studies in both rodents and primates shows that mICCs are organized in a continuum rather than clusters and homogenous in terms of cell types [19, 21-24]. Thus, we also cannot rule out that modulation of BMA PACAPergic inputs to different sub-divisions of the mICCs lead to similar effects on fear behaviors. However, we did not find sex differences in PACAPergic manipulation from BMA to mICCs.

Deletion of PAC1 receptors from mICCs led to sexually dimorphic effects on contextual fear behaviors. In females, PAC1 deletion decreased fear acquisition, without altering fear generalization or extinction. PAC1 receptors in mICCs of females could modulate the asymptote of the learning curve. In males, PAC1 receptors in the mICCs appear to be important for regulating fear generalization and extinction. Deletion of PAC1 receptors from mICCs in males enhanced fear generalization and reduced extinction of fear. This finding reflects that in males PAC1 receptors in this anatomical juncture are necessary for regulating fear in an inappropriate context, whilst ablating them leads to enhanced fear. The reduced extinction also indicates a failure to inhibit acquired fear when the PAC1 receptors are lacking. Overall decreased fear levels were observed in females and heightened levels of fear were observed in males when PAC1 receptors are absent in mICCs. PAC1-contaning mICC neurons have projections to various regions besides the CN [19, 25]. In our studies, we found that fear behaviors can be differentially impacted when PACAP versus its receptor PAC1 are manipulated in the mICCs. One obvious reason is that the neuropeptide, PACAP and its receptors are expressed in different neurons. The fact that we saw inconsistencies in sex differences in fear behaviors only when the receptor is manipulated indicates that there may be differences in neurons that express these receptors that project to other brain regions. Taken together, the behavioral findings indicate that sensory and associative information are differentially regulated by PACAP and PAC1 via the BMA-mICC microcircuit.

Amygdala microcircuitry is heterogeneous and complex in its morphology and function. Fear-related sensory information is thought to flow from the BLA complex to CN either directly or via mICCs, which in turn alters fear expression via direct or indirect projections to the periaqueductal grey. Each region in this circuit can also independently modulate fear [26-28]. In particular, functions of the mICCs are influenced by a variety of neuropeptides. Our finding that PACAP and PAC1 influences different aspects of fear behaviors indicates that PACAP/PAC1 is a critical neuropeptide system in this anatomical juncture. Holistically, our findings underscore the importance of mICCs, which despite being tiny structures in the interface of BLA/BMA and CN can regulate several aspects of contextual fear behaviors.

Previous anatomical studies suggested that BMA projections to mICCs are sparse. In those studies, phytohemagglutinin lectin anterograde labeling method was used. While this method of labeling provides useful information about regional association, it is limited in providing information about transneuronal connections, defining functional synapses or providing information about genetically-defined cell populations [29, 30]. Using two complementary approaches we found that the BMA sends heavy PACAPergic projections to dorsal and ventral mICCs.

*Ex vivo* optogenetic stimulation of the PACAPergic projections from BMA to the mICCs led to enhanced EPSCs with a short latency (10 ms) in mICCs neurons. This indicates that these projections are most likely monosynaptic. We also showed that the morphological properties of mICCs neurons are similar to medium spiny neurons and are highly excitable, in agreement with previous findings regarding these neurons and their high input resistance [22-24, 31]. We were able to abolish the EPSCs with CNQX, indicating that these responses are mediated by glutamatergic AMPA receptors. However, it is possible that NMDA receptors play a role. NMDA receptors mediate calcium entry in response to synaptic stimulation and have been previously shown to modulate mICC function [17, 32, 33]. In a sub-population of mICCs neurons, we observed an EPSC followed by a longer latency IPSC, which could be due to collateral projections from nearby mICC neurons. We cannot rule out the possibility that the IPSCs occurred due to signal backpropagation along the dendrite. This in turn could activate other bifurcated projections to mICCs leading to IPSCs via an alternate pathway within this region. Previous studies have shown that action potentials in the mICCs can back-propagate but are modulated by voltage-dependent potassium channels. The functions of these channels are mediated by neuromodulators and second messenger systems [34]. We used a PAC1 receptor antagonist concurrently with optogenetic stimulation and perhaps surprisingly found that the EPSCs were further enhanced. This suggests that PACAP is co-released with glutamate from PACAPergic neurons. It is also possible that PACAP 6-38 blocked the basal tone of PACAP and not the co-release. Release of PACAP may mitigate the postsynaptic influence of glutamate when these neurons are stimulated by shunting inhibition via PACAP-dependent opening of potassium channels. While PAC1 receptors are virtually absent in the BMA, it is not known whether the PAC1 receptors are present on the neuronal terminals originating from the BMA in form of auto receptors. If these receptors are in fact present on the BMA neurons innervating the mICCs, then they would be in a perfect position to modulate the observed physiological properties via presynaptic mechanism of modulating neuropeptide release.

While fear is a natural response that keeps organisms safe when faced with danger, dysregulation of fear such as in PTSD can seriously cripple an individual’s ability to function. A traumatic stress results in acquisition of a conditional fear at a greater magnitude, over-generalization of fear to an otherwise safe context, enhanced recall of fear and decreased extinction of fear. We looked at all these aspects of fear in our study. One theory of heightened fear in PTSD is due to inappropriate inhibitory control over fear, leading even mild reminders of trauma to trigger strong symptoms and reduced propensity to extinguish acquired fear [35-42]. PACAP and PAC1 have previously been shown to be important for fear regulation. High blood levels of PACAP, especially in females, and methylation of PAC1 in a sex-independent manner were previously associated with PTSD [11]. In rodents, PACAP has been shown to enhance consolidation and extinction of contextual fear and enhance excitatory synaptic transmission in projections from BLA to lateral division of CN [43-46]. Given the role of PACAP and PAC1 in contextual fear, we chose to delineate the role of the PACAP and PAC1 in the BMA and mICCs microcircuitry in regulation of different aspects of contextual fear behaviors in both male and female mice. While mICC neuronal architecture is complex, these neurons do appear to be plastic in their response to sensory information and modulate fear behaviors in a heterogenous manner that is dependent upon the information coming into this region [15, 17, 47, 48].

Our findings are important because mICCs are major cell groups in the amygdala microcircuitry and process specific aspects of fear-related information [10, 27, 49]. Studying the role of microcircuit like BMA-mICCs in fear regulation via a specific neuropeptide system like PACAP/PAC1 provides better anatomical knowledge regarding the substrates that can be harnessed for targeted therapies for ameliorating symptoms in disorders like PTSD [37, 50, 51]. Hence in the future it will be of interest to determine if PACAP interacts with other neurotransmitter or neuropeptide systems to regulate fear behaviors [18].

## Methods

### Experimental Models and Subject Details

All experimental procedures were conducted in accordance with the guidelines set by the National Institute of Health and the Institutional Animal Care and Use Committee at the University of California, Los Angeles. All mice were kept on *ad libitum* access to food and water in a light- and temperature-controlled vivarium. Mice (3–4 months) were housed in plastic clear cages (3-5 mice/cage as littermates) in the vivarium with lights on at 7 AM and lights off at 7 PM. Experiments were performed between 9 AM and 3 PM.

#### Mouse Lines

Three different mouse lines were used in this study. The first was a Tg (Adcyap1-EGFP) FB22Gsat/Mmucd (RRID:IMSR_MMRRC:012011) reporter mouse line that faithfully expresses enhanced green fluorescent protein in PACAP positive neurons. These mice were generated using the bacterial artificial chromosome (BAC; RP24-358O1) by the Gene Expression Nervous System Atlas (GENSAT) project and obtained from the Mutant Mouse Resource and Research Centers. These mice were backcrossed from FVB/NTac to C57BL/6 for at least five generations [12]. The second mouse line was an Adcyap1r1^loxP/loxP^ mouse. These mice were generated in a C57BL/6 background with a conditional knockout (KO) allele (PAC1^loxP/loxP^ mice) through the NIH-funded knockout mouse project (KOMP). The third mouse line was an Adcyap1-2A-Cre mouse line. These mice (Adcyap1-2A-Cre) target Cre to most populations of PACAP neurons of the brain including the amygdala [10] (Allen Brain Atlas).

#### Measure of freezing

Freezing is a complete lack of movement except for respiration [52]. Freezing was measured using VideoFreeze (Med-Associates Inc.) that performed real-time video recordings at 18 frames per second. With this program, adjacent frames are compared to provide the grayscale change for each pixel and the sum of pixels changing from one frame to the next constitutes a momentary activity score. To account for video noise and to approximate scoring by a trained human observer a threshold is set at 18 activity units so that an instance of freezing is counted when that the activity score remains below this threshold for 1 sec [53]. Percentage freezing=Freezing Time /Total Time×100 for a period of interest. Data are presented as mean percentages (+/- SEM).

Because Med-Associates software uses the number of pixels changed across the entire video frame to calculate the amount of freezing, we were not able to use it for automated analysis of freezing for our optogenetic studies. This is because even when the mice were freezing, the movement in the optogenetic cable was calculated by the software as movement. Hence for optogenetic behavioral experiments, we analyzed freezing using ezTrack software [54], which enables removal of this cable artifact. In brief, videos were cropped to reduce the influence of optogenetic cables in the upper portion of the field of view. Subsequently, the number of pixels whose grayscale value changed from one frame to the next was calculated. Freezing was then scored when this number dropped below an experimenter-defined threshold for at least 30 frames (1 second). The freezing threshold was determined by visual inspection of the video and by comparing a subset of the results obtained to the results of manual scoring. All cropping and thresholding parameters were identical across sessions. Data are presented as mean percentages (+/- SEM).

### Determining BMA to mICCs PACAPergic Innervation

#### Immunofluorescence for visualizing expression of PACAP-EGFP neurons

For immunofluorescence labeling, 40-micron coronal brain sections were cut from (Adcyap1-EGFP) mice (N=5; M=2, F=3). Sections were blocked and permeabilized in a solution containing PBS + 10% normal goat serum (NGS) + 1% bovine serum albumin (BSA) + 0.5% TritonX-100 for 1 hour. An anti-GFP primary antibody (A11122, Life Technologies) and anti-NeuN antibody (MAB377, Millipore) were diluted 1:500 in PBS + 5% NGS + 1% BSA and sections were incubated overnight at 4°C, then washed in PBS, and subsequently incubated with an anti-rabbit AlexaFluor 488 (Life Technologies Cat#A11008, RRID:AB_10563748) and anti-mouse Cy3 (Abcam Cat#Ab97035, RRID:AB_10680176) secondary antibodies diluted 1:400 for 2-4 hours at room temperature. Sections were washed in PBS and mounted on slides, and coverslipped with Prolong Gold Antifade Reagent (Life Technologies). Fluorescence images were acquired with a Keyence widefield microscope (BZ-X710).

#### Intersectional viral method for labeling PACAPergic neurons from BMA to mICCs

To further validate the existence of PACAPergic projections from BMA to mICCs, we utilized an intersectional viral labeling method. For this, we used an hEF1α-LS1L-mCherry-IRES-flpo virus (Harvard Vector Core), with either AAV5-EF1a-fDIO-ChR2-eYFP-WPRE or AAV5-EF1a-fDIO-eYFP-WPRE (UNC Vector Core) viruses. Adcyap1-2A-Cre mice (N=3) went through stereotaxic surgeries with BMA co-ordinates: L/M: +/− 3.25; A/P: −1.7; D/V: −4.75) and mICCs co-ordinates: -L/M: +/− 2.7; A/P: −1.06; D/V: −4.2. We first microinfused hEF1α-LS1L-mCherry-IRES-flpo into the mICCs and the AAV5-EF1a-DIO-hChR2(H134R)-mCherry virus into the BMA. The hEF1α-LS1L-mCherry-IRES-flpo virus is a (Cre) recombinase dependent and retrogradely transporting virus, so when injected into the mICCs of Adcyap1-2A-Cre mice, it only expresses in PACAP-containing neurons as they express Cre in the mICCs terminal where it is injected. This virus in turn contains flippase (FLP) recombinase in its sequence. So, we then microinjected a FLP-dependent AAV5-EF1a-fDIO-ChR2-eYFP-WPRE or AAV5-EF1a-fDIO-eYFP-WPRE virus into the BMA. This allows the ChR2 or eYFP (control) to be expressed specifically in PACAPergic neurons that project from the BMA to mICCs. The volume of viral injections in mICCs was 0.1-μl over 2 minutes and in the BMA was 0.3 μl over 6 minutes and 10 minutes of diffusion time for each infusion. Following the surgical procedure, in all experiments, mice were allowed to recover for 21 days to allow viral transduction.

#### Visualizing expression of PACAPergic neurons

After recovery from surgery, mice were euthanized, brains extracted, and 40-micron brain sections were cut. Sections were mounted on slides and coverslipped with Prolong Gold Antifade Reagent (Life Technologies). Fluorescence images were acquired with a Keyence imager.

### *In vivo* Optogenetic Stimulation of BMA-mICCs PACAPergic Neurons and Fear Behaviors

#### Intersectional viral method for expressing ChR2 specifically in PACAPergic neurons

We wanted to determine if altering the activity of PACAPergic neurons that innervate the mICCs changes fear behavior. A previous study showed that the ex vivo optogenetic method can be used to analyze and study the role of intercalated cells in regulating fear behaviors, so we utilized optogenetics to answer our question [16]. For these experiments, we used the Adcyap1-2A-Cre mice and carried out the same intersectional virus labeling strategy described previously. Briefly, we injected the Cre-dependent hEF1α-LS1L-mCherry-IRES-flpo in the mICCs and AAV5-EF1a-fDIO-ChR2-eYFP-WPRE or AAV5-EF1a-fDIO-eYFP-WPRE in the BMA for specifically expressing ChR2 in PACAPergic neurons that project from BMA to ICCs. Two 200 µm diameter optic fibers were also implanted bilaterally above mICCs or BMA (available from Prizmatix) and cut at the length of 4.6 mm or 5 mm, respectively. The core diameter of the fibers was 250um and an outer diameter of 275 µm. The numerical aperture (NA) was 0.66. For the stimulation, the optical fiber was connected to an LED emitting blue light (473 nm, 20 Hz train of 10 ms on/off pulses, 10 min duration). Optogenetic stimulation was carried out bilaterally in each mouse throughout the duration of the experiment.

#### Behavioral Procedure

Conditioning Apparatus: Mice were run individually in sound and light attenuated conditioning boxes (Med Associates Inc., Georgia, VT) (Fig. 5). The boxes were equipped with Near Infra-Red (NIR) Video Fear Conditioning System and could be configured to represent different contexts by changing the internal structure, floor texture, illumination, and odor. Context A (28 × 21 × 21 cm) had a clear Plexiglas back wall, ceiling, and front door with aluminum sidewalls. It also had a grid floor with evenly spaced stainless-steel rods cleaned and was scented with 50% Windex. The floor in context A was connected to a scrambled foot shock generator. Context B had a clear Plexiglas back wall, ceiling, and door with aluminum sidewalls. The chamber was altered by adding a white curved sidewall that extended across the back wall. The floor of context B consisted of an acrylic white board floor, cleaned and scented with 1 % acetic acid solution. The room was illuminated with red light and the visible overhead lights were turned off in the box.

#### Behavioral design

For each behavior day, mice were lightly restrained, the optic cable was connected to the indwelling fiber optics in their head and placed in the testing chamber. We tested acquisition, generalization, and extinction of fear as measured by freezing.

Acquisition: For acquisition we placed the animals in a context for four minutes and thirty seconds every day around the same time for 5 days. At the fourth minute each day, the mice received a 1 second .65 mA shock. After 29 seconds they were removed from the chamber and put back in their home cages, where they were housed with littermates. Mice were transported to the laboratory together in their home cages. There was one rest day between acquisition and generalization testing.

Generalization Test: The animals were placed in a completely different context (B) for four minutes and thirty seconds. No shocks were delivered.

Retention Test: One day after generalization testing the animals went through fear retention tests in context A. Mice were placed in the context for 4 minute 30 seconds without any shocks.

Extinction Test: For the next 5 days animals went through fear extinction, again in context A. Extinction sessions were 30 minutes long. We measured freezing for the first 4 minutes, but the animals remained in the chamber for the entire 30 minutes.

### Deletion of PAC1 Receptors From the mICCs and Measurement of Fear-related Behavior

For experiments involving deletion of PAC1 receptors, we microinfused AAV2-hsyn-GFP-Cre or AAV2-hsyn-GFP into mICCs using the stereotaxic co-ordinates L/M: +/− 2.7; A/P: - 1.06; D/V: −4.2. Although it is challenging to precisely target small structures like the medial ICCs, we have shown feasibility of confining virus infusion to such a small structure by using a specialized digital stereotax (Model 1900, David Kopf Instruments) and pulled glass pipettes that are commonly used for electrophysiological recordings with single cell resolution. We verified through various methods that we were able to precisely target the mICCs, including DAPI infusion (Fig. S2C), infusion of AAV expressing mcherry (Fig. S2D), infusion of AAV2-hsyn-GFP (Fig. S2E), and then co-labeling INTRESECT virus with an antibody against FoxP2, protein that is highly expressed in the mICCs (Fig. S3C). Using these methods, we found that viral infusions were constrained by the surrounding capsule if the placement was accurate. The behavioral procedure and design were identical to the optogenetic experiments. The only difference was for measuring freezing, automated Medassociates Videofreeze software was used as described previously.

#### Validation of PAC1 receptor deletion using dual *in situ* hybridization with RNAscope

After the behavioral trials were complete the mice were sacrificed, and their brains extracted and immediately stored in at −80 degrees celsius. The brains were sliced at 15 microns in a cryostat and slices containing the amygdala were collected on microscope slides. The deletion of PAC1 receptors was verified using RNAscope for analyzing expression of RNA tissue sections (ACD Biotechne). Briefly, we performed *in situ* hybridization steps following RNAscope^®^2.5 HD Duplex Assay protocol for fresh frozen sections. After completion of the labeling, sections were cover-slipped using Prolong Gold (Thermo Fisher Scientific) with 4’,6-diamidino-2-phenylindole (DAPI) and the edges were sealed with clear nail polish. PAC1 mRNA puncta counts were carried out in 4 serial sections containing the mICCs that started at the same rostral plane that were captured with a 40X objective. mICCs frame was 25 microns x 25 microns size. Analysis of dots was conducted using FIJI software. For each image DAPI+PAC1 mRNA and DAPI+GFP were analyzed separately. For counts, DAPI positive cells and PAC1 RNA dots were counted separately and expressed as mean grain count.

### Measurement of cfos Expression After *in vivo* Optogenetic Stimulation of BMA-PACAP Neurons that Innervate the mICCs

#### Viral surgeries and *in vivo* optogenetic stimulation

For these experiments, we utilized the same intersectional approach as described previously for labeling PACAPergic neurons projecting from BMA to mICCs by infusing hEF1α-LS1L-mCherry-IRES-flpo into the mICCs and AAV5-EF1a-fDIO-ChR2-eYFP-WPRE or AAV5-EF1a-fDIO-eYFP-WPRE into the BMA. A 200 µm diameter optic fiber was also implanted bilaterally above BMA (available from Prizmatix) and cut at the length of 5 mm. The core diameter of the fibers was 250um and the outer diameter was 275 µm. The numerical aperture (NA) was 0.66. After 21 days, mice were anesthetized with isoflurane and the optical fiber was connected to a LED emitting blue light (473 nm, 20 Hz train of 10ms on/off pulses, 10 min duration). Optogenetic stimulation was carried out unilaterally and the hemisphere of stimulation was counter-balanced between mice. Ninety minutes following the simulation, mice were sacrificed, and brains extracted, cryoprotected and frozen. During the ninety minutes, the mice were in their home cages without anesthesia.

#### Immunohistochemistry for measuring cfos expression

The brains were processed for cfos immunohistochemistry. First, 40-micrometer coronal sections containing the amygdala were collected serially. On day 1, tissue sections were washed in 1×TBS three times for five minutes, then blocked in 1mL of 1×TBS with 5% Normal Donkey Serum, 0.1% BSA and 0.3% Triton-X for 1 hour. Then the tissue sections were incubated overnight at 4 degrees Celsius with the primary goat polyclonal to cfos (1:500, 24 h, abcam; RRID: SCR_012931) primary antibody. According to the manufacturer, this antibody is a ‘synthetic peptide conjugated to Blue Carrier Protein by a Cysteine residue linker corresponding to the internal sequence amino acids 283–295 of Human c-Fos (<u>NP_005243.1</u>). On the second day, the sections were washed in 1×TBS three times five minutes each and then incubated in the Alexa 488 donkey anti-goat secondary antibody (1:200, Life Technologies) for 2 hours at room temperature. After washing with 1×TBS for 3 times 5 minutes each, tissue sections were mounted on glass slides and cover-slipped using Prolong Gold (Thermo Fisher Scientific) with 4’,6-diamidino-2-phenylindole (DAPI) and the edges were sealed with clear nail polish. Positive cfos immunolabeling was analyzed and quantified in brain sections containing the mICCs.

#### Cell counts for cfos experiments

The numbers of cfos+ cells were manually counted in the region of the mICCs by two trained researchers that were blind to the treatment conditions. Separately cfos+ cells co-expressed with chR2 expressing fibers were also manually counted and the percentage of chR2 /PACAP cells expressing cfos+ over only cfos+ was calculated.

### Electrophysiological recordings of mICCs neuronal activity and *ex vivo* optogenetic stimulation of PACAPergic neurons that innervate mICCs

#### Electrophysiology and *ex vivo* optogenetics

For *ex vivo* electrophysiology experiments, we nfused in the BMA of the Adcyap1-2A-Cre mice a Cre-dependent AAV5-EF1a-DIO-hChR2(H134R)-mCherry virus (UNC Vector Core) (N=6; M=3, F=3). After a 21-day recovery from surgery, animals were deeply anesthetized with isoflurane and decapitated amygdala slices were prepared. The brains were placed in ice-cold modified artificial cerebrospinal fluid (aCSF, containing, in mM: 194 sucrose, 30 NaCl, 4.5 KCl, 1 MgCl_2_, 26 NaHCO_3_,1.2 NaH_2_PO_4_, and 10 D-glucose) and cut into 300 μm-thick coronal slices containing the intercalated cell (ICC) layer of the amygdala. The slices were then allowed to equilibrate for 30 min at 32–34°C in normal aCSF (containing in mM; 124 NaCl, 4.5 KCl, 2 CaCl_2_, 1 MgCl_2_, 26 NaHCO_3_, 1.2 NaH_2_PO_4_, and 10 D-glucose) continuously bubbled with a mixture of 95% O_2_/5% CO_2_, stored at room temperature in the same buffer, and used for experiments within 6 hr of slice preparation.

Electrophysiological methods were described previously [55, 56]. Cells were visualized with infrared optics on an upright microscope (BX61WI, Olympus). pCLAMP10 software and a MultiClamp 700B amplifier was used for electrophysiology (Molecular Devices). For these recordings the intracellular solution in the patch pipette contained the following, in mM: 135 potassium gluconate, 3 KCl, 0.1 CaCl_2_,10 HEPES, 1 EGTA, 8 Na_2_-phosphocreatine, 4 Mg-ATP, 0.3 Na_2_-GTP, pH 7.3 adjusted with KOH and filtered with a 0.2 µm syringe filter. For biocytin labeling, 2 mg/ml biocytin (Tocris) was dissolved in the intracellular solution and cells were dialyzed for 20 minutes. For all recordings, the patch-pipette tip resistance was ∼5 MΩ. The initial access resistance was less than 25 MΩ for all cells and if this increased by more than 5 MΩ the cell was discarded.

We identified mICCs by their somatic morphology and location in the dorsal ICC nucleus, elevated membrane resistance and the presence of a slowly accommodating inward current upon hyperpolarization, in agreement with past work [15]. The concentration of drugs applied onto brain slices via bath perfusion were: 10 μM CNQX (Cayman chemical), 250 nM PACAP 6-38 (Tocris), 20 µM Bicuculline (Cayman chemical), and 0.5 μM Tetrodotoxin (Cayman chemical). All compounds were stored at −20°C as stock solutions and diluted in ACSF just prior to use. ChR2-mediated responses were evoked by 470 nm light flashes from an LED source (Sutter Instrument) at power of 0.025 mW/mm^2^ and for a flash duration of 25 ms each.

#### Viruses

For the intersectional method, we used hEF1α-LS1L-mCherry-IRES-flpo (MIT and Harvard Vector Core) and AAV5-EF1a-fDIO-ChR2-eYFP-WPRE or AAV5-EF1a-fDIO-eYFP-WPRE (UNC Vector Core) constructed under Dr. Rachel Neve and Dr. Karl Deisseroth, respectively. For *ex vivo* electrophysiology experiments, we used Cre-dependent AAV5-EF1a-DIO-hChR2(H134R)-mCherry virus (UNC Vector Core). For deletion of PAC1 receptors we used the AAV 2-hsyn-GFP-Cre or AAV2-hsyn-GFP (UNC vector core).

### Microscopy for All Experiments

The tissue sections were analyzed using a Keyence BZ-X700 -All-in-One Fluorescence Microscope. Images were analyzed with Fiji image processing software (NIH, Bethesda, MD; RRID: SCR_002285). Images were converted to binary mode (black and white image). For *ex vivo* studies, a confocal microscope was used for analyzing the expression of biocytin-filled cells in the mICCs.

### Statistical Analysis for All Experiments

To be consistent, for every phase of testing (days 1-14), we measured freezing for the first 4 minutes of the session. This corresponds to the preshock period on the acquisition days, providing a measure of contextual fear that is not confounded by the unconditional behavioral effects of the shock. For the behavioral experiments in which acquisition and extinction were measured, a three-way analysis of variance (ANOVA) was used to measure differences in means with two between (sex and group) and one within (day) factors. For the other behavioral experiments, a two-way ANOVA was carried out to measure differences in the means with group and sex as between group factors. Significant effects indicated by the ANOVA were further analyzed with a post-hoc Bonferroni post-hoc analysis. The level of significance used for all analyses was p < 0.05. For electrophysiological recordings, non-parametric Mann-Whitney U tests were performed.

## Acknowledgements

AKR is now at Icahn School of Medicine at Mount Sinai, New York, 10029, USA. We would like to thank Dr. Avishek Adhikari for valuable input for the optogenetic experiments in this project. These experiments were supported by NRSA-F32 MH10721201A1-AKR, NARSAD 26612-AKR, R21MH098506 (MSF/JW MPI), RO1MH062122 (MSF) and the Staglin Center for Brain and Behavioral Health (MSF).

## Author Contributions Section

AKR designed the studies, ran experiments, and wrote the manuscript. JCO ran electrophysiological experiments. ZP helped with optogenetic behavioral experiment analysis. SG and JT helped with optogenetic experiments. JC, NE and NK helped with behavioral experiments. WZH provided the Adcyap1-2A-Cre mice. RLN made the HSV viruses used in the INTRSECT approach. JW made and provided the PAC1^loxP/loxP^ and the Adcyap1-EGFP mice. BK helped with all aspects of electrophysiological experiments in his lab and provided feedback. MSF supervised all the studies and helped design the experiments and manuscript preparation.

## Figure Legends

**Fig. S1. PACAP-expressing neurons in the BMA innervate the medial ICCs**

**A.** Representative image panels showing expression of PACAP-EGFP and DAPI in Adcyap1-EGFP mice.

**B.** Image panels showing expression of EGFP in Adcyap1-EGFP mice in the BMA and with terminals in the mICCs.

**C.** PACAP mRNA expression in the BMA and mICCs compared to PACAP-EGFP expression. PACAP mRNA and EGFP expression is high in the BMA.

**Fig S2: *Expression levels of PAC1 mRNA and examples of site-specific infusions in the mICCs***

**A.** *PAC1 mRNA expression in the mICCs from Allen Brain atlas.*

**B.** *In situ hybridization for PAC1 mRNA using antisense and sense probes against PAC. PAC1 mRNA is high and specific in the mICCs.*

**C.** *Infusion of DAPI into the mICCs to verify placement in the mICCs.*

**D.** *Infusion of AAV-CHR2 in the mICCs to verify viral expression and site-specific targeting of mICCs.*

**E.** *Example PAC1 mRNA expression in the amygdala and mICCs with and without AAV expressing Cre-GFP. AAV injections were restricted to the capsular region of the mICCs.*

**Fig S3: *Example images of PAC1 expression***

***A.*** *and* ***B***. *RNAscope in situ* hybridization labeling with control probes.

**C.** Verification of expression of FoxP2 (marker of mICCs) co-localized with ChR2 in the mICCs.

**D.** Graphs depicting fear retention test after deletion of PAC1 receptors from the mICCs. There was no effect of Sex or Group or Group x Sex interaction on the retention test (F=2.463; p>0.05).

**Fig S4: Representative electrophysiological traces of mICCs neurons after stimulation of PACAP BMA fibers**

**A.** Representative current traces (*left*) from ICCs in response to trains of blue light flashes with or without CNQX present and average EPSC amplitude (*right*) at each flash for a train of 40 flashes.

**B.** Representative current trace (*left*) from ICCs in response to single flashes with or without bicuculline applied. Average IPSC amplitude (*right*) in response to single flashes with or without bicuculline.

**Fig S5: Representative images of biocytin-filled neurons in mICCs after electrophysiological recordings**

**A.** Example image showing a biocytin-filled neuron in the mICCs at the site of recording

**B.** Examples of image reconstruction of two neurons in the mICCs filled with biocytin after electrophysiological recordings

