## Supplemental Figures for "A peptidergic amygdala microcircuit modulates sexually dimorphic contextual fear"

### Slide 1
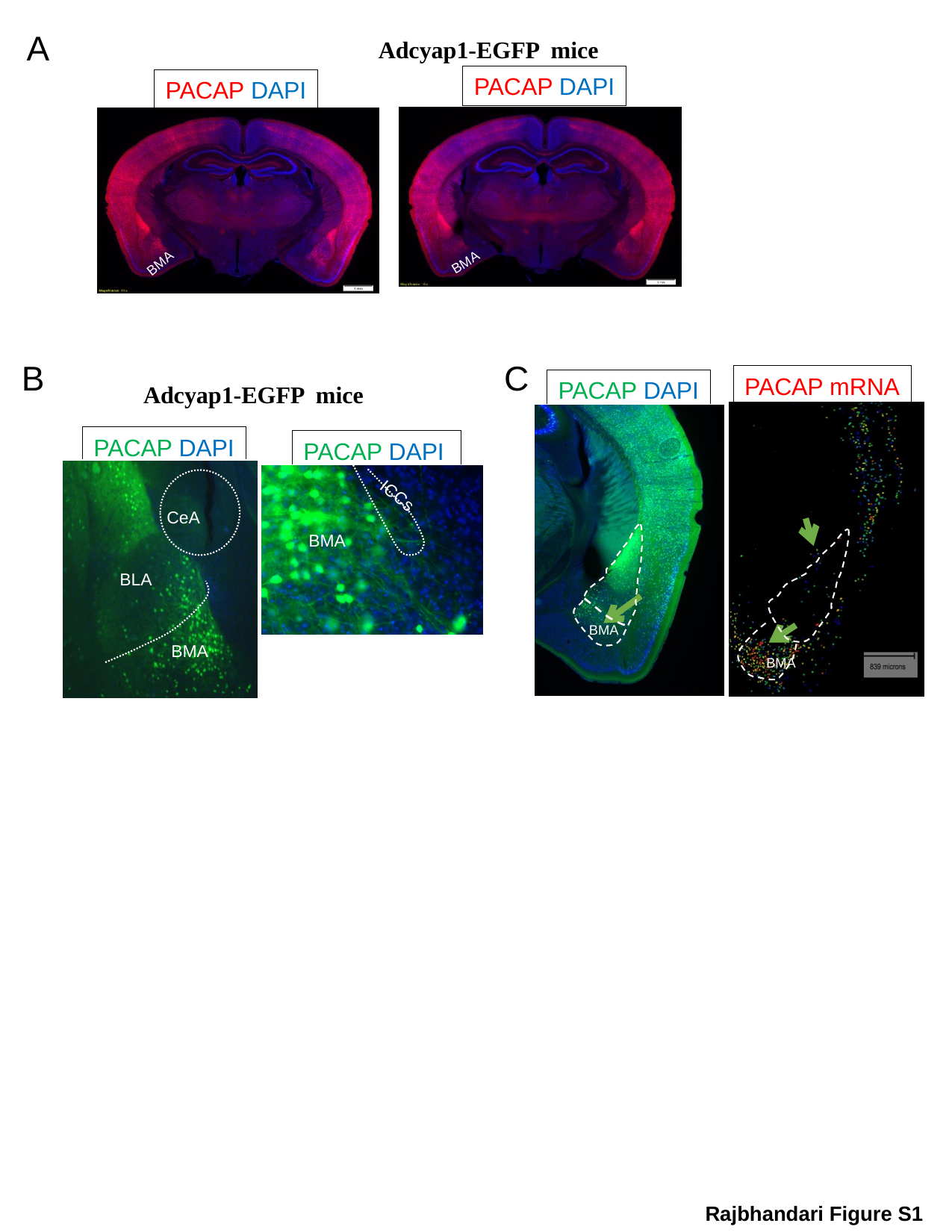

A
Adcyap1-EGFP mice
PACAP DAPI
BMA
PACAP DAPI
BMA
B
C
PACAP mRNA
PACAP DAPI
Adcyap1-EGFP mice
BMA
PACAP DAPI
PACAP DAPI
ICCs
BMA
CeA
BLA
BMA
BMA
BMA
Rajbhandari Figure S1

### Slide 2
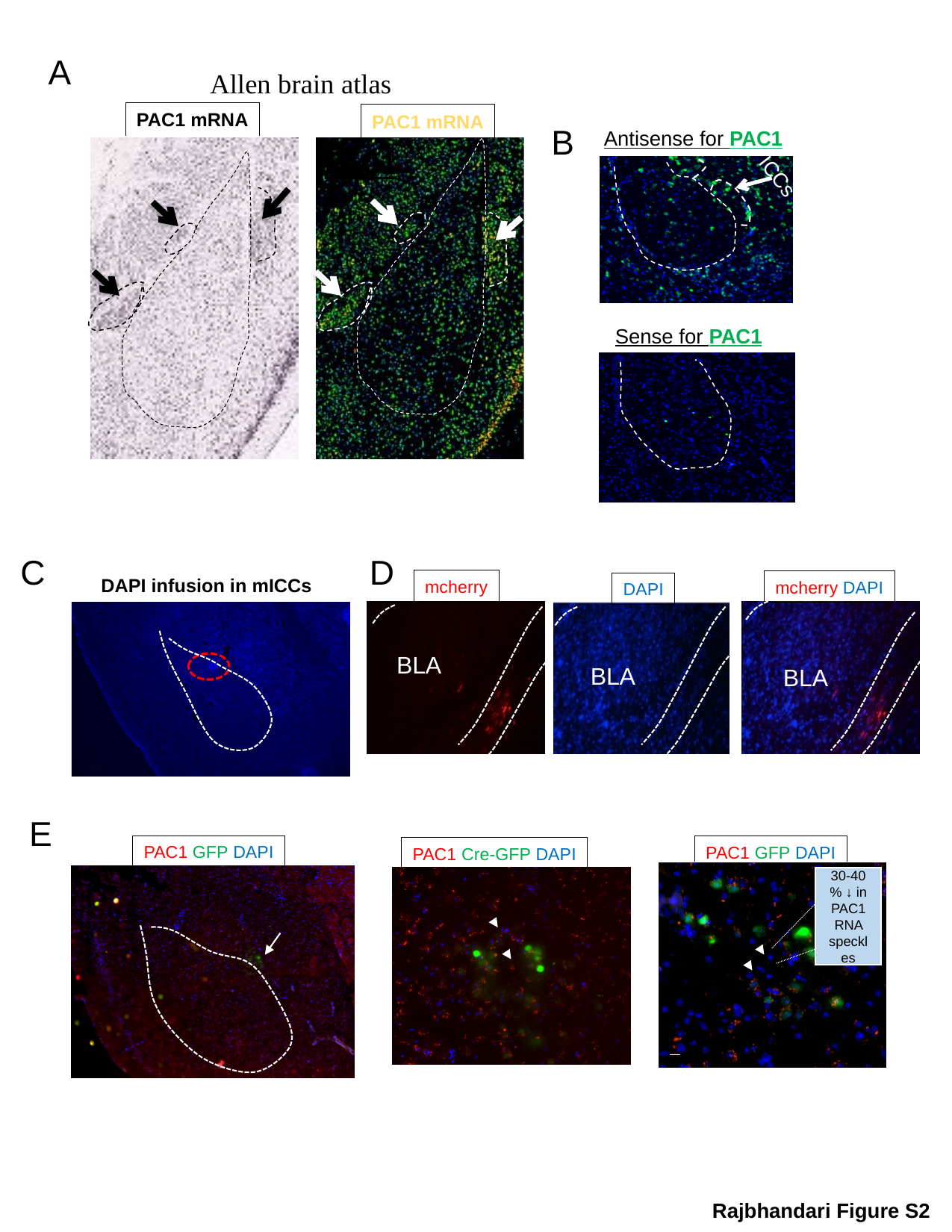

A
Allen brain atlas
PAC1
PAC1 mRNA
PAC1 mRNA
B
Antisense for PAC1
ICCs
Sense for PAC1
C
D
DAPI infusion in mICCs
mcherry
mcherry DAPI
DAPI
BLA
BLA
BLA
E
PAC1 GFP DAPI
PAC1 GFP DAPI
PAC1 Cre-GFP DAPI
30-40 % ↓ in PAC1 RNA speckles
AAV-Cre-GFP
Control
AAV-GFP
Rajbhandari Figure S2

### Slide 3
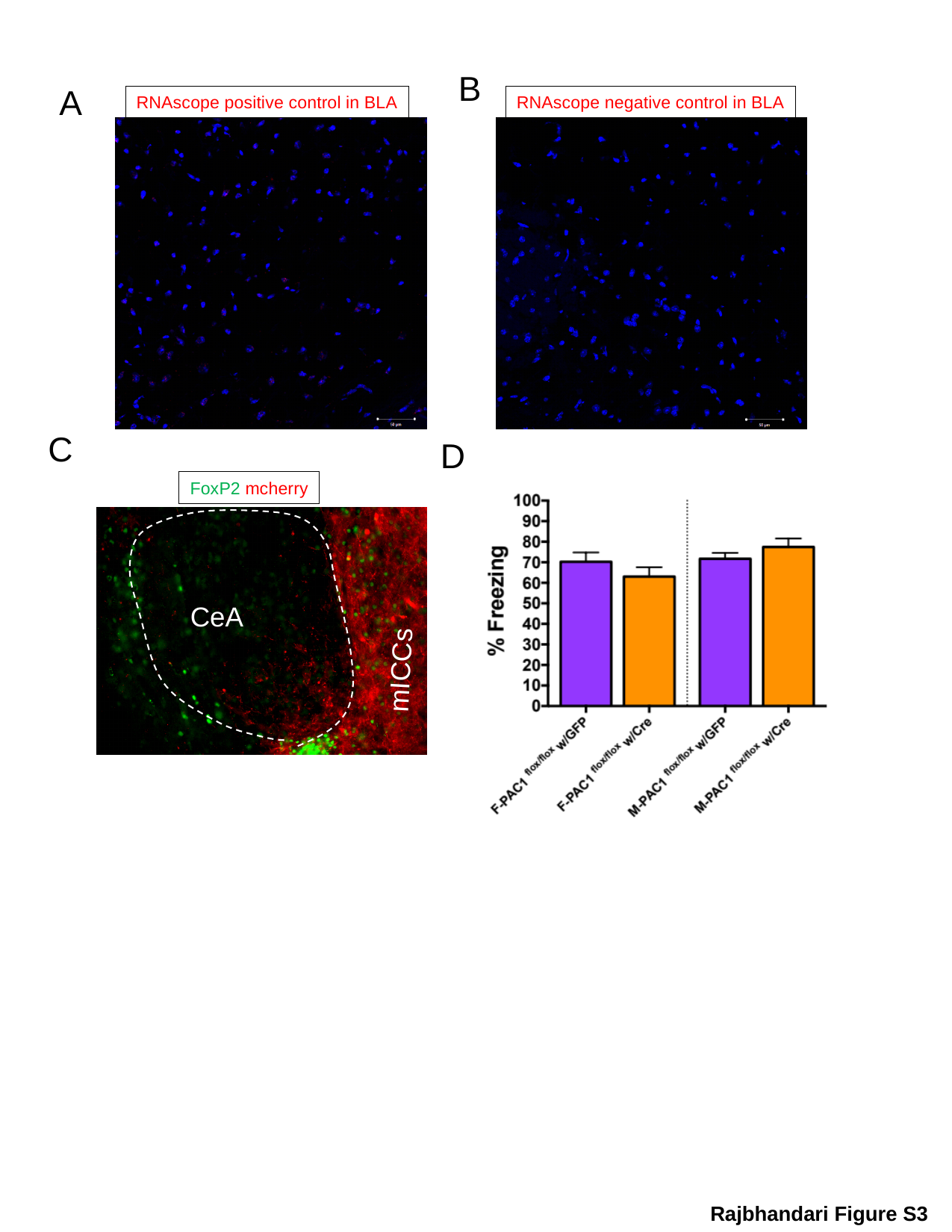

B
A
RNAscope negative control in BLA
RNAscope positive control in BLA
C
D
FoxP2 mcherry
CeA
mICCs
Rajbhandari Figure S3

### Slide 4
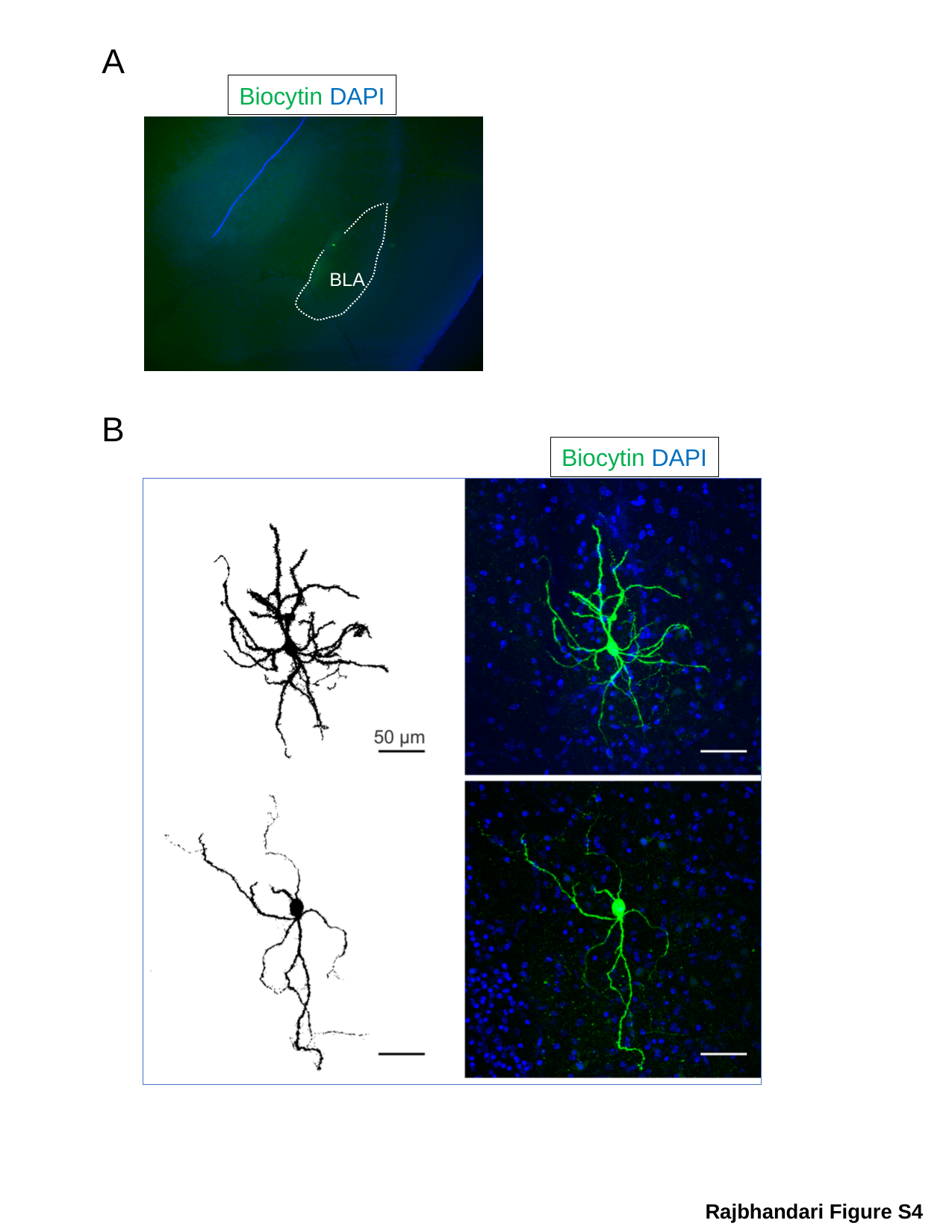

A
Biocytin DAPI
BLA
B
Biocytin DAPI
Rajbhandari Figure S4

### Slide 5
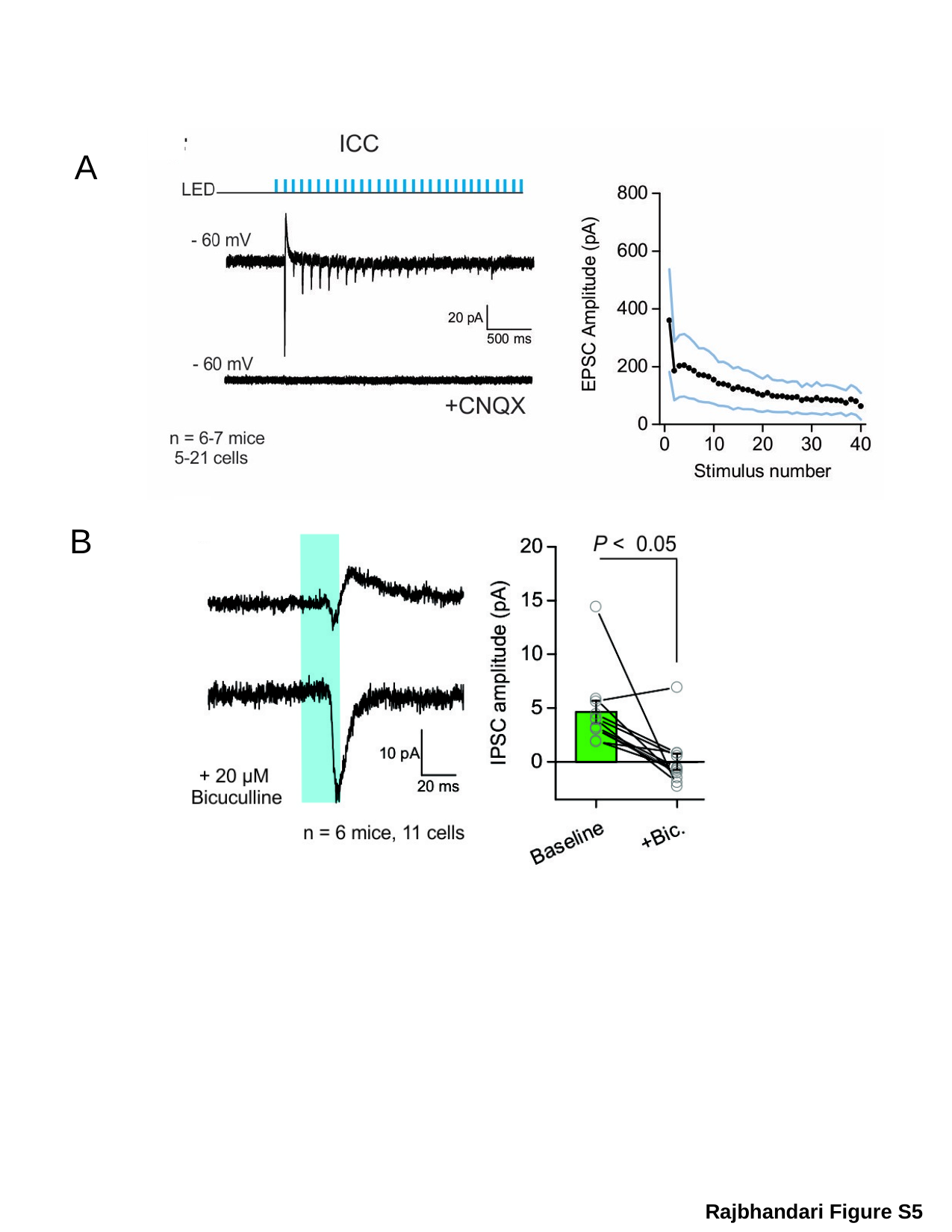

A
B
Rajbhandari Figure S5
